# VRK3 depletion induces cell cycle arrest and metabolic reprogramming of Pontine Diffuse Midline Glioma (DMG)-K27 altered cells

**DOI:** 10.1101/2023.03.24.533960

**Authors:** Virginie Menez, Thomas Kergrohen, Tal Shasha, Claudia Silva-Evangelista, Ludivine Ledret, Lucie Auffret, Chloé Subecz, Manon Lancien, Yassine Ajlil, Kévin Beccaria, Thomas Blauwblomme, Estelle Oberlin, Jacques Grill, David Castel, Marie-Anne Debily

## Abstract

We previously identified VRK3 as a specific vulnerability in DMG-H3K27M cells in a synthetic lethality screen targeting the whole kinome. The aim of the present study was to elucidate the mechanisms by which VRK3 depletion impact DMG-K27M cell fitness. Gene expression studies after *VRK3* knockdown emphasized the inhibition of genes involved in G1/S transition of the cell cycle resulting in growth arrest in G1. Additionally, a massive modulation of genes involved in chromosome segregation was observed, concomitantly with a reduction in the level of phosphorylation of serine 10 and serine 28 of histone H3 supporting the regulation of chromatin condensation during cell division. This last effect could be partly due to a concomitant decrease of the chromatin kinase VRK1 in DMG following *VRK3* knock-down. Furthermore, a metabolic switch specific to VRK3 function was observed towards increased oxidative phosphorylation without change in mitochondria content, that we hypothesized would represent a cell rescue mechanism. This study further explored the vulnerability of DMG-H3K27M cells to VRK3 depletion suggesting potential therapeutic combinations, *e.g.* with the mitochondrial ClpP protease activator ONC201.

## Introduction

Diffuse midline glioma (DMG) H3K27-altered represents the most severe pediatric brain tumor. Almost all the affected children die within 2 years following diagnosis^1, 2^ without significant improvement of their overall survival in the last decades despite numerous trials to improve the outcome by adding new agents to the radiotherapy standard ^3^. These tumors harbor a global loss of K27 trimethylation of histone H3 resulting from the substitution of a lysine residue to a methionine (K27M) in the regulatory tail of histone H3 ^4–6^ in 90% of cases. This mutation occurs almost exclusively in *H3-3A*, encoding H3.3 histone variant or in *H3C1*, encoding H3.1 variant ^4^ and is considered the initiating driver event of DMG oncogenesis leading to a broad dysregulation of the epigenome. However, this alteration could not be specifically targeted until now. At present, only secondary driver events are targetable which limits the efficacy of these treatments. Therefore, we performed a synthetic lethality screen by RNA interference targeting the human kinome to unravel genes required for pontine DMG-K27M cell survival. We identified *VRK3* (Vaccinia related kinase 3) as an essential gene that has not yet been associated with DMG oncogenesis ^7^.

VRK3 belongs to the Vaccinia Related Kinase (VRK) family containing 3 distinct members in mammals: VRK1, VRK2 and VRK3 serine/threonine kinases, which are mainly located in the nucleus. VRKs are known to regulate several aspects in cell such as cell cycle progression, nuclear envelop dynamic during mitosis and apoptosis ^8, 9^. These proteins also play a role in histone post-translational modifications (PTM) ^10, 11^. In particular VRK1, the most studied member of the VRK protein family, has been shown to bind to chromatin either during mitosis ^8^ or during DNA damage response ^12^. VRK1 is in particular required for chromatin compaction in G2/M and during mitosis it progressively regulates histone H3 through its interplay with AURKB, by phosphorylating threonine 3 and serine 10 (Ser10 or S10) residues sequentially. The role of VRK2 has been less investigated, but was associated with a downregulation of apoptosis ^13^. VRK3 was recently classified as a pseudokinase and remains poorly studied in comparison with other VRK members.

All VRK kinases were shown to phosphorylate BAF a regulator of post-mitotic nuclear envelope formation ^14, 15^. The partial overlap of VRK1- and VRK3-interacting proteins reflects their common involvement in cell cycle regulation, chromatin assembly and DNA repair ^16^ while exhibiting distinct roles in these processes. Both VRK1 and VRK3 were shown to promote liver cancer progression, VRK1 regulating G1/S transition and mitosis and VRK3 S phase progression ^16^. VRK1 was also shown to regulate G1/S transition in fibroblasts ^17^ but was recently associated with G2/M arrest in glioblastoma ^18^, suggesting a cell-dependent impact on cell cycle. At a clinical level, *VRK1* overexpression has been associated with poor prognosis in many solid tumors including high-grade glioma ^19, 20^. We observed that in adult glioma high *VRK3* expression was correlated with decrease in overall survival independently of the grade of the tumor ^7^, whereas *VRK2* expression was associated with an increase of survival in high-grade astrocytoma ^21^.

This study aims to improve our knowledge on VRK3 function and the molecular impact of its repression in the context of DMG-H3K27 altered. We confirmed that *VRK3* repression leads to a strong and rapid arrest of cell growth, resulting in a blockage of the cells in G1 phase and also emphasize a global dysregulation of DMG metabolism.

## Results

### Impact of *VRK3* knockdown on DMG, H3K27M mutant transcriptome

In order to investigate possible mechanisms underlying the observed phenotype of cell fitness decline following decrease of *VRK3* in pontine DMG-K27 altered cells ^7^, we examined differences in global gene expression. RNA-seq was performed 44h and 60h post-transduction with two distinct shRNAs targeting *VRK3* to be able to evaluate the early impact of *VRK3* knockdown (KD) in four independent *in vitro* models of DMG. *VRK3* expression presented a fold reduction ranging from 1.53 to 7.89 (mean=3.17) at 44 hours, and from 1.33 to 8.25 (mean=3.42) at 60 hours (FigS1A), with GSC2 and GSC3 harboring the highest KD.

The most important source of variation in the data after correction of individual biological variability correspond to the modulation of *VRK3* expression as reflected by PCA analysis based on the whole transcriptome data. Indeed, all samples with *VRK3* KD are well separated from a second group containing both non-transduced cells and cells transduced with non-targeting control shRNAs (Fig1A). Sample unsupervised clustering supports this observation with a clear separation of samples transduced with shVRK3 from the others (Fig1B). Samples with *VRK3* KD were almost perfectly subdivided into two subgroups according to the shRNA targeting *VRK3* used (shVRK3-1 or shVRK3-4). This can be explained by the greater KD with shVRK3-1 compared to shVRK3-4 (FigS1A), even though both shRNAs were designed to target the same set of alternative transcripts of *VRK3*.

**Fig1.**
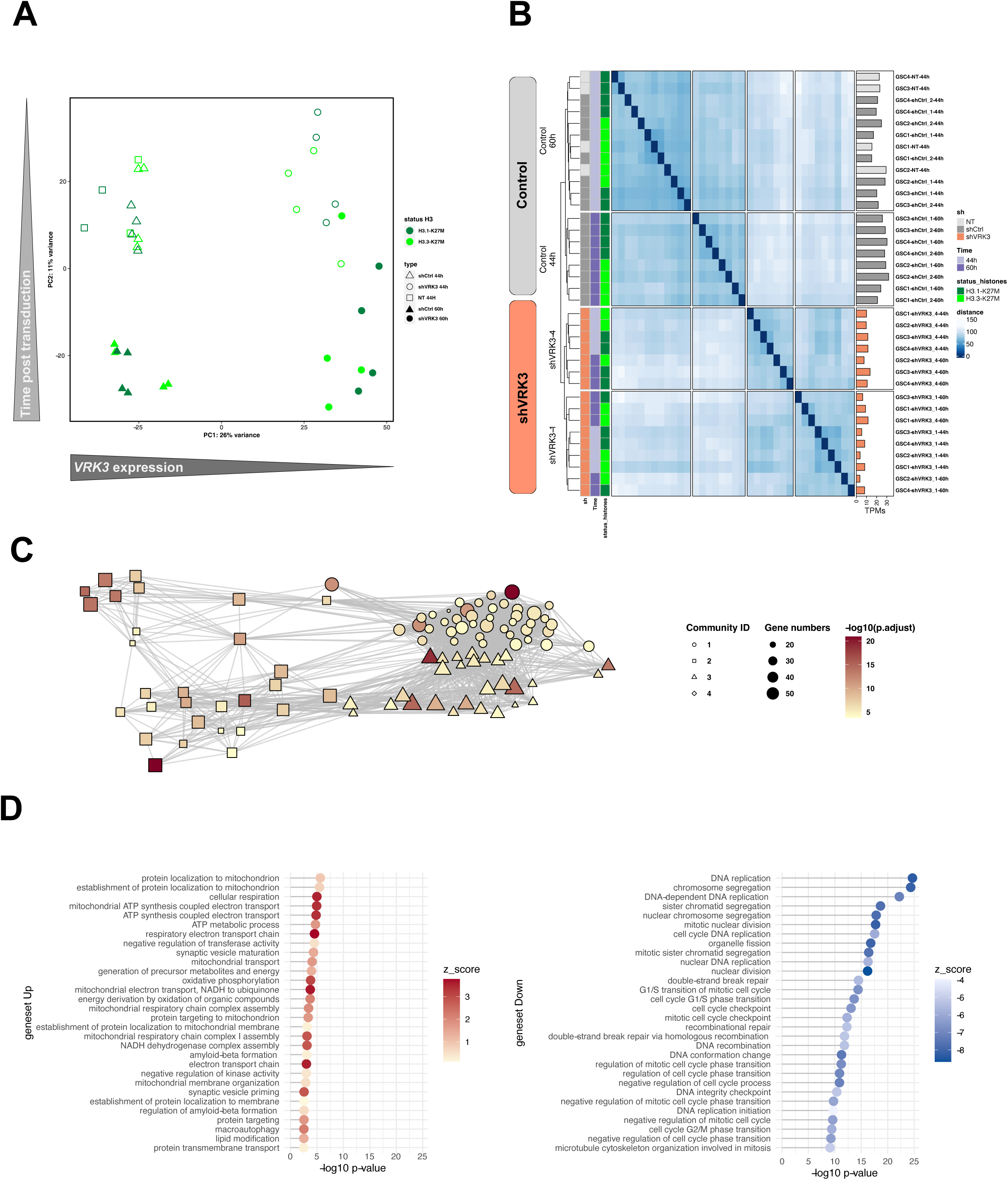
Main pathways affected by *VRK3* repression in DMG H3K27M. A-PCA analysis of samples based on the entire RNAseq dataset (32 334 genes). B-Sample unsupervised clustering based on the whole transcriptome. The dendrogram illustrates the ordering of patient samples in 2 clusters corresponding to samples transduced with shVRK3 or control samples (not transduced NT or transduced with shCTRL1 or shCTRL2). C – Enrichment map from the top 100 genesets. Relationships between genesets as depicted by Enrich distill analysis. The size of the node encodes the information of the number of DEGs while the color is representative of the computed Z score for each set. The color is indicative of the significance of expression changes (antilog of pvalue) of genes assigned to a particular set. The distinct communities among the enrichment map were identified by Louvain clustering. D – Geneset enrichment analysis of DEG associated with adjusted pvalue threshold of 0.001. The 30 most enriched genesets among upregulated (red) and downregulated genes (blue) are presented, showing *e.g.* significance and direction of change (z-score)

Samples are spread on PC2 according to time post-transduction but similarly with negative control shRNA and shRNA targeting *VRK3* reflecting the impact of transduction on *in vitro* DMG models (Fig1A). Accordingly, shCTRL samples at 44- or 60-hours post-transduction are distributed in distinct subclusters in consensus clustering (Fig1B). However, the comparison of modulated genes following *VRK3* KD at both time points 44- and 60-hours post-transduction exhibits an elevated correlation (Pearson correlation 0.64, FigS1B), and higher than the comparison of modulated genes according to their H3 mutational status (Pearson correlation 0.43, FigS1C). Yet, differential analysis between shVRK3 cells 44- and 60-hours post-transduction revealed a few modulated genes (20 upregulated and 32 downregulated) in comparison with the contrast shVRK3 *versus* shCTRL (1264 upregulated and 1626 downregulated) with the same threshold of adjusted pvalue ≤ 0.001 (FigS1D&E), suggesting that most effect is already present at 44h. Functional annotation of differentially expressed genes (DEGs) in shVRK3 cells between the 2 timepoints showed an enrichment of genes involved in histone acetylation with overexpression of *HDAC5* (FC 1.56; padj 0.003), *HDAC8* (FC 1.44; padj 0.0005), *TRERF1* (FC 1.54; padj 0.00002) and downregulation of *RBM14* (FC 0.64; padj 5.64 e-07); and cholesterol biosynthetic process with downregulation of *DHCR7* (FC 0.53; padj 0.0002), *NSDHL* (FC 0.59; padj 0.0003), *FASN* (FC 0.74; padj 0.0024), *ACAT2* (FC 0.066; padj 0.003) and *NFYA* (FC 0.76; padj 0.007).

Consequently, samples with *VRK3* KD were grouped together for the majority of further differential expression analysis.

Geneontology enrichment analysis of DEGs after *VRK3* KD pinpointed a network of the top 100 genesets, from which four communities of highly connected genesets were identified by Louvain clustering (Fig1C). Three of them, associated with ‘kinetochore and chromatin compaction’, ‘regulation of cell cycle transition’, ‘telomere organization’ present multiple connections with each other’s (Fig1C, FigS2 & Table S2). The fourth appears completely independent and is labelled as ‘protein localization in mitochondrion’.

Independent functional annotation identified a larger number of functionally enriched biological processes among downregulated genes, in accordance with the volcano plot of differential analysis results showing more downregulated DEGs (FigS1D&E, Table S3). A massive repression of genes involved in distinct phases of the cell cycle was observed, reflecting that the main impact of *VRK3* KD is a cell growth arrest (Fig1D) whereas the top 30 categories of upregulated genes emphasize an impact on mitochondrial metabolism.

We did not observe major differences between H3.1- and H3.3-mutated cells, as shown by the overlap among the top 20 modulated genes of each subgroup following *VRK3* decrease (Fig2A&D), except that higher fold changes are present in H3.1-K27M GSC reflecting most likely some heterogeneity between the two H3.3-K27M GSC models (Fig2B-D and FigS1C). In accordance, the same biological processes appeared enriched after *VRK3* repression in the two subgroups even though chromosomal segregation and G1/S transition are impacted to a greater extent in H3.1-K27M cells (Fig2A).

**Fig2.**
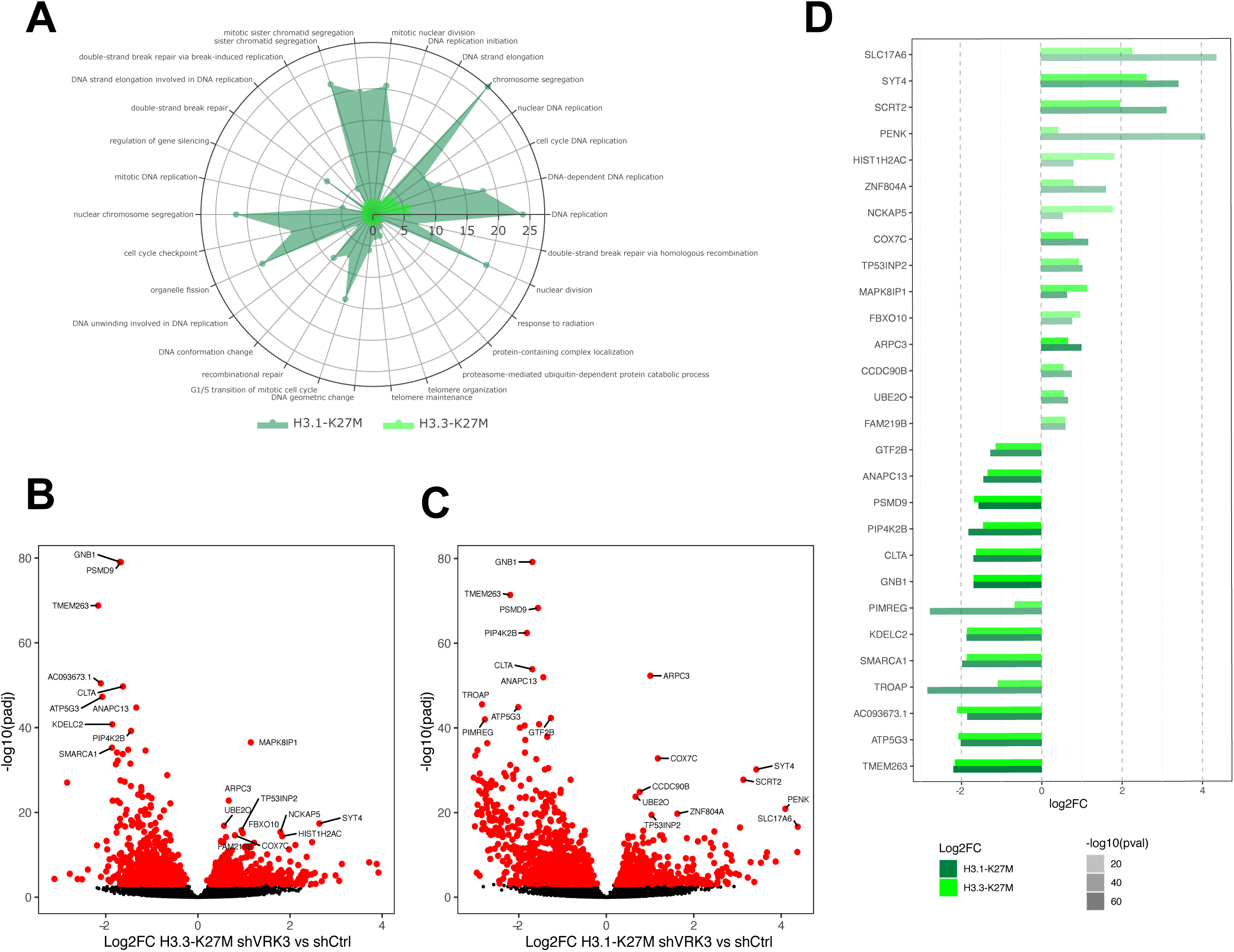
Similar impact of *VRK3* KD in H3.1-K27M and H3.3-K27M cells. A. Radar plot displaying the 30 most enriched genesets in H3.3 GSC following *VRK3* KD ranked by decreasing antilog of pvalue (res_enrich_H33, light green). The corresponding pvalue of overrepresentation analysis in H3.1-mutated tumors are presented (res_enrich_H31, dark green). Functional analysis performed by enrichGO using biological process gene annotation. B. Volcano plot of gene expression in H3.3 shVRK3 cells *versus* H3.3 shCTRL cells plotting antilog of adjusted pvalue on y-axis *versus* log2 Fold Change on x-axis. DEGs associated with adjusted pvalue <0.001 are color-coded in red. The 10 most upregulated and downregulated genes are indicated. C. Volcano plot of gene expression in H3.1 shVRK3 cells *versus* H3.1 shCTRL cells plotting antilog of adjusted pvalue on y-axis *versus* log2 Fold Change on x-axis. DEGs associated with adjusted pvalue <0.001 are color-coded in red. The 10 most upregulated and downregulated genes are indicated. D. Bar charts presenting the expression of genes belonging to the top ten upregulated and downregulated genes in H3.1 (light green) and/or H3.3-mutated cells (dark green) after *VRK3* KD. Gene modulation is presented as the log2FC of RNAseq data on x-axis and the significance is encoded by color transparency.

### *VRK3* KD disturbs kinetochore function and chromosome segregation

RNAseq data emphasized a strong impairment of mitosis and more specifically chromosome segregation one of the two main functions impacted following *VRK3* KD (Fig1D, Fig2A and Fig3A). Among the top modulated genes, we found a considerable downregulation of clathrin (*CLTA*) which is involved in kinetochore stabilization (FC 0.1; padj 1.4 e-106) and a significant overexpression of *ARPC3* (FC 3.19; padj 5.4 e-075), encoding a member of the ARP2/3 complex involved in actin polymerization and regulating the dynamics of the actin cytoskeleton in particular during cell division.

**Fig3.**
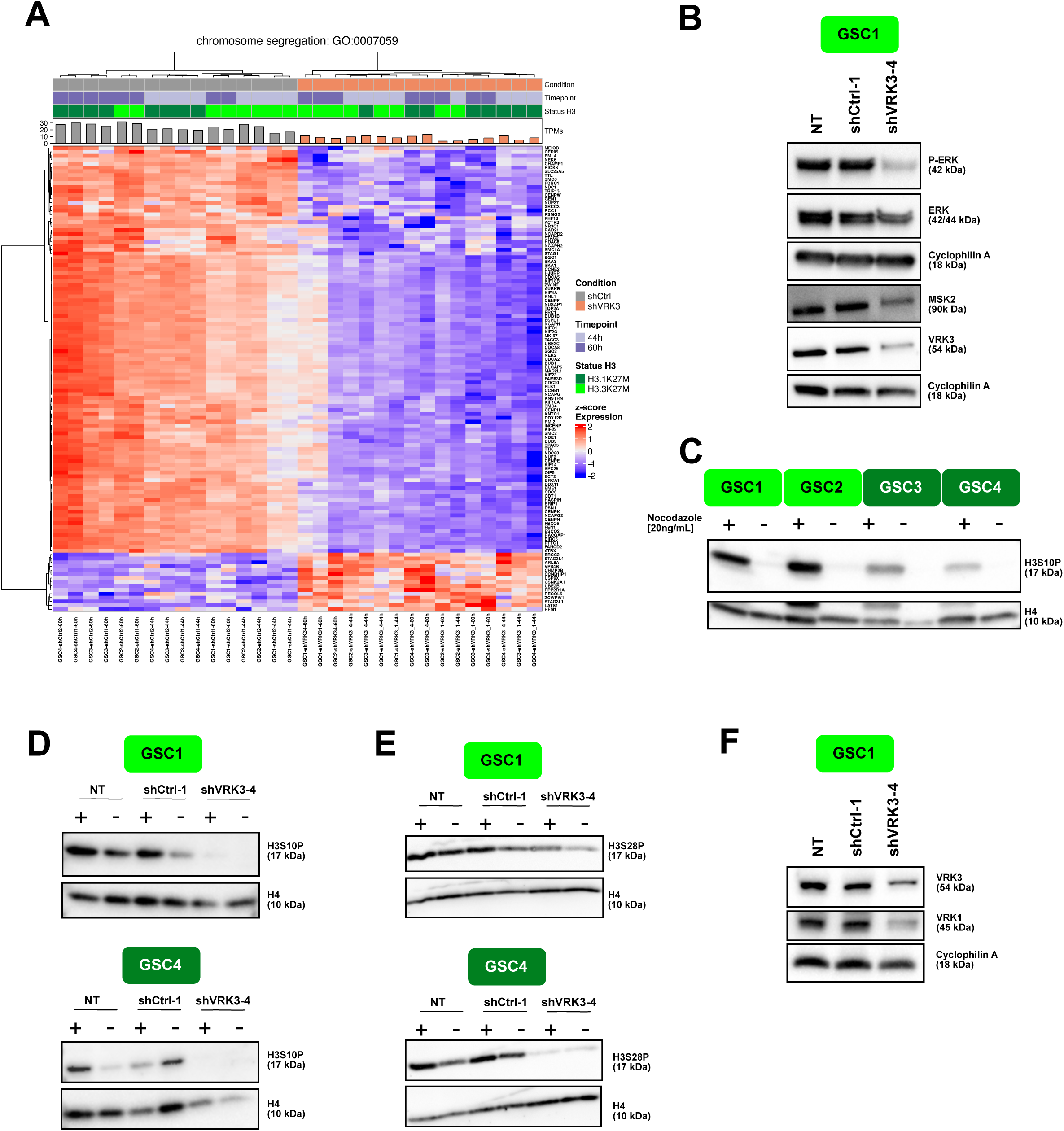
*VRK3* impair chromosome segregation/affect chromatin dynamics in DMG. A-Hierarchical clustering and heatmap of modulated genes after *VRK3* KD involved in chromosome segregation (Gene Ontology GO:0044843). B- Protein modulation of VRK3, ERK and MSK2 96 h after transduction with shCTRL-1 or shVRK3-1. 4. Cyclophilin was used as loading control. C- Western Blot analysis showing levels of H3S10P in GSC1, GSC2 (H3.3-K27 model) and GSC3, GSC4 (H3.1-K27M) with or without nocodazole treatment. Histone H4 was used as loading control. D-E- Western Blot analysis showing levels of H3S10P and H3S28P in H3.3-K27M GSC1 and H3.1-K27M GSC4 with or without nocodazole treatment 168h after transduction with shCTRL-1 or shVRK3-4. Histone H4 was used as loading control. F-Protein modulation of VRK1 and VRK3 168h after transduction in GSC1. Cyclophilin was used as loading control.

VRK3 was previously shown to affect cell proliferation by negatively regulating ERK activity in distinct cell types ^10^. We evaluated the impact of *VRK3* depletion in DMG cells and observed opposite results. Indeed, ERK was slightly decreased at protein level, while its phosphorylation was almost fully abolished (Fig3B).

This inhibition of phospho-ERK could consequently affect several mitotic kinases operating downstream such as mitogen and stress activated protein kinase MSK1 and MSK2. Among these, MSK2 protein content was decreased in DMG after *VRK3* KD as shown by western blot in GSC1 (Fig3B) and a reduction of phospho-MSK2 was also observed in phospho-array experiments performed in GSC2 and GSC4 cells (FigS3A).

Interestingly, in a previous lethality screen, we found that shRNAs targeting *MSK2* negatively impacted proliferation of DMG cells, but to a lesser degree than *VRK3* KD^7^. We evaluated the impact of pharmacological inhibition of MSK2 using two multi-targeted tyrosine kinase inhibitors. One of them did not present any efficiency in both GSC and NSC cells (SB747651A, IC50 ranging from 15 to 61.3 uM, *data not shown*). NSC4 appeared more sensitive than H3.3-mutated cells to the second drug, R0318220 which is not concordant with the results of lethality screen (FigS3B). This discrepancy presumably results from a lack of specificity of the drug and/or suggests that the strong impact of *VRK3* inhibition in DMG cannot be solely mimicked by MSK2 signaling inhibition.

MSK2 is a canonical H3S10 kinase. MSK2-dependent H3S10 phosphorylation plays a main role in chromatin dynamics and this post-translational modification is also tightly associated with cell cycle progression. H3S10 phosphorylation (H3S10P) increases as cells enter mitosis, with a high occupancy of H3S10P nucleosomes along the entire length of chromosomal arms during prophase and metaphase ^22^. Few H3S10P-positive cells were detected in basal conditions in DMG cells, even though this percentage was higher in H3.3-K27M than in H3.1-K27M GSCs and normal NSCs (FigS3C). These differences remained observable in western blot after a nocodazole-block in M-phase to increase the global content of H3S10P (Fig3C and FigS3D). Also, *VRK3* KD reduced the pool of H3S10P-positive cells (Fig3D and FigS3E).

In addition to MSK2, other key mitotic regulators of H3S10P mark deposition were modulated after *VRK3* repression. *AURKA* (FC 0.16; padj 1.27E-11) localizes at centrosomes and mitotic spindle poles throughout mitosis and is involved in centrosome maturation and separation. *AURKB* (FC 0.06; padj 6.72E-25) localized to the kinetochore and the anaphase central spindle is expressed in early mitosis and phosphorylates H3S10 to aid at chromosome condensation and segregation.

H3S10P is involved in two structurally opposed processes; *i.e.* in addition to its involvement in chromosome compaction during cell division, this PTM is regulated during transcription activation^23^. Indeed, during interphase, H3S10 phosphorylation at certain promoters leads to chromatin remodeling during transcription^22^. MSK2 also induces phosphorylation of both ser10 and ser28 of histone H3 when cells are exposed to certain environmental stresses, giving access to transcription factors in regulatory DNA sequences. Similarly to H3S10P, the proportion of H3S28P-positive cells is higher in GSC than in NSC (FigS3F), in particular in H3.3-mutated cells, and decreases following *VRK3* KD (Fig3F & FigS3G).

A previous study of the VRK3 interactome highlighted VRK3 interactions with several chromatin components like histone H1, H2A, H2B type 1 & 2, H3.3, H4 or epigenetic regulators such as Jumonji protein (JARID2), suggesting a role for VRK3 in chromatin organization ^16^. We found among VRK3 substrates a significant enrichment for genes involved in telomere organization (FigS3H). Also, 24 genes belonging to VRK3 interactome (n=108) were differentially expressed in shVRK3 cells (FigS3I). Among them, *JARID2* encoding a core subunit of PRC2 complex was slightly upregulated (FC 1.56; padj 0.008) and *HIST1H2AC* (encoding histone H2A) belonged to the top upregulated genes following *VRK3* KD in our study (FC 6.27; padj 2.02E-16) (Fig2D).

Moreover, a significant overlap of VRK3 and VRK1 interactomes was reported (40 common interacting proteins) as well as their direct binding to each other ^16^. Among the common substrates of VRK1 and VRK3, four were significantly modulated with a threshold of adjusted pvalue of 0.001 after *VRK3* KD: three ribosomal proteins (RPL11 - FC 1.41, RPS14 FC 1.35 and RPS15A FC 1.5) and NPM1 exhibiting several functions including H3, H4 and H2B core histone chaperoning function (FC 0.49). Interestingly, chromatin condensation was reported to be enhanced via phosphorylation of histone H3 on Ser10 by VRK1 ^24^. VRK1 harbors mainly a nuclear localization in DMG with a significant proportion bound to chromatin as reported in others cellular contexts ^17^(FigS3J). In comparison, VRK3 was also mostly found in the nucleus but mainly in the soluble nuclear fraction (FigS3K), although we detected the presence of a small proportion of VRK3 protein in the cytoplasm by immunofluorescence, suggesting a potential association with organelles (*data not shown*). Also, VRK2 protein localization was shown to display a similar cytoplasmic localization by confocal microscopy, indicating a binding to endoplasmic reticulum and to a lesser extend to mitochondria ^25^. Strikingly, we observed a decrease of VRK1 protein following *VRK3* KD by RNA interference, even greater than VRK3 itself (Fig3F). The level of expression of both genes was correlated in DMG primary tumors contrary to *VRK2* and *VRK3* (FigS3L). Thus, the modulation of H3S10 phosphorylation observed in our cells after *VRK3* KD could result, at least partly, from VRK1 decrease and not only from a direct effect of *VRK3* KD.

### *VRK3* KD impairs DMG proliferation by a massive repression of genes involved in the G1/S cell cycle transition

Overall, *VRK3* inhibition led to G1 growth arrest as reflected by the significant increase of the percentage of G1 cells and loss of cells in S phase 96h after transduction (Fig4A&B). We observed some variations between GSC models in the distribution of cells among the distinct phases of cell cycle even without shVRK3 transduction. GSC1 appeared less sensitive than the others to shVRK3, possibly linked to a lower *VRK3* KD in these cells (Fig4A&B, FigS1B). We did not observe considerable variations in *VRK3* expression level during the cell cycle in untransduced cells. Yet, an increase was measured in G2 phase both at transcriptional (from 1.28 to 1.86, pvalue<0.01, FigS4A) and protein level (FigS4B).

**Fig4.**
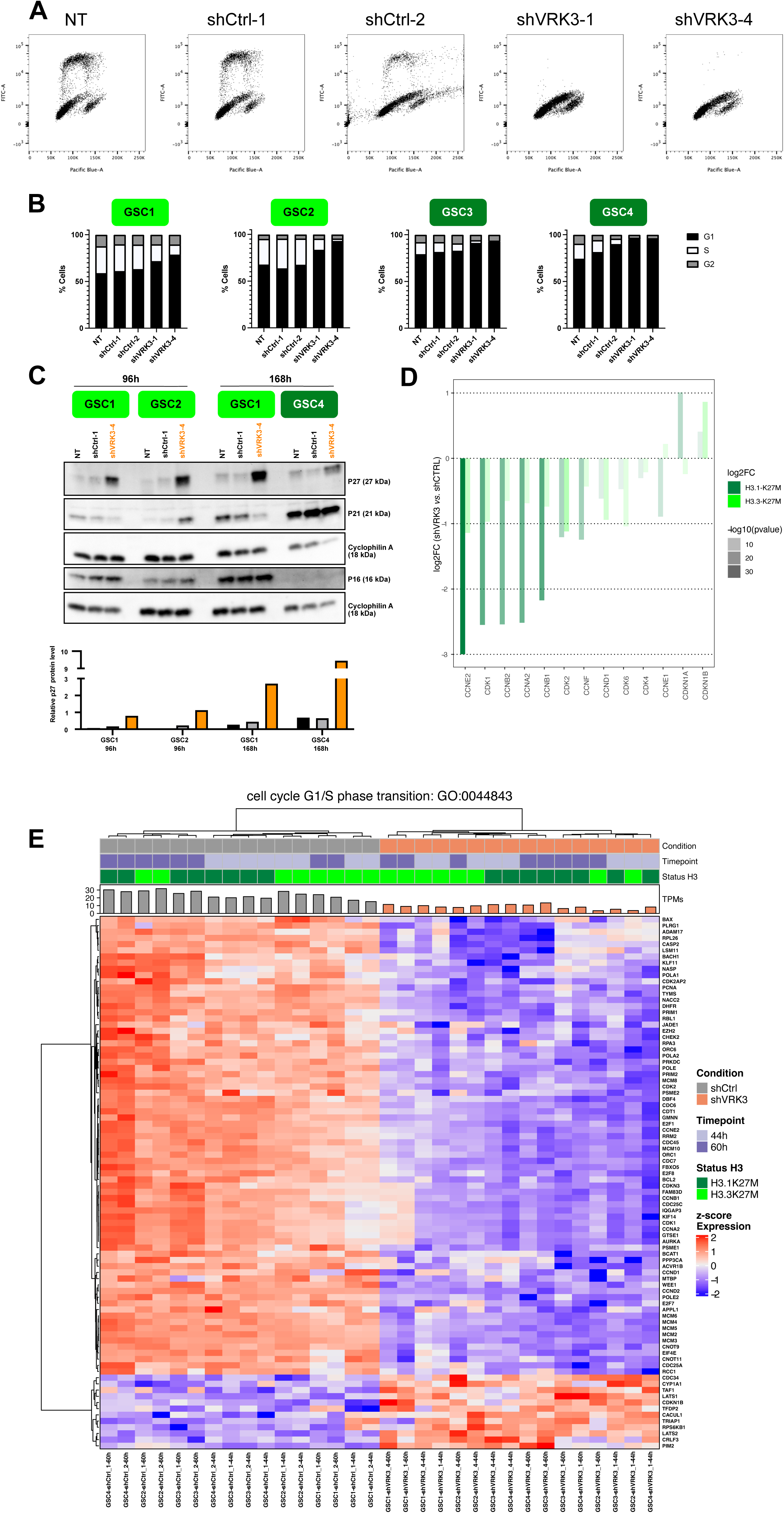
*VRK3* inhibition led to DMG-K27altered G1 growth cell arrest. A-Two-dimensional plots (EdU *vs.* FxCycle Violet stain) of GSC4 cells 96h hours after transduction with shCtrl-1, shCtrl-2, shVRK3-1 and shVRK3-4 and quantification of the different cell-cycle population. At least thirty thousand cells were recorded for each condition, triplicates were performed for each GSC model excepted duplicates for GSC3 cells. B- Histogram presenting percentage of the different cell-cycle populations in GSC cells (H3.3-K27M models GSC1, GSC2 in light green, H3.1-K27M models GSC3, GSC4 in dark green) 96h after transduction at M.O.I. 3 with shCTRL-1, shCTRL-2, shVRK3-1 and shVRK3-4. At least thirty thousand cells were recorded for each condition, quadriplicates were performed for each GSC model excepted duplicates for GSC3 cells. C- Hierarchical clustering of DEGs in shVRK3 *versus* shCTRL belonging to cell cycle G1/S phase transition geneset (Gene Ontology GO:0007059). Heatmap show RNAseq expression level (Z-score on normalize expression matrix) D-Bar charts presenting modulation of CDK and CDK inhibitors in H3.3 and H3.1-mutated cells after *VRK3* KD. Gene modulation is presented as the log2FC of RNAseq data on y-axis and the significance is encoded by color transparency. E-Western Blot analysis showing levels of P27, P21 and P16 in GSC1 and GSC2 (H3.3-K27 model) and GSC4 (H3.1-K27M) 96h or 168h after transduction with shCtrl-1 or shVRK3-4. Cyclophilin was used as loading control. Densimetric quantification of western blot was processed with Image Lab and P27 protein level was normalized to cyclophilin expression.

In accordance with cell cycle modifications, RNAseq data showed a significant modulation of genes involved in cell cycle G1/S transition (Fig1D and Fig4C) as early as 44h hours after transduction and to a greater extent in H3.1-mutated cells (Fig2A and Fig4D). All cyclins and CDKs were downregulated, with a huge repression of cyclin E2 (*CCNE2*) and *CDK2* in both H3.1 and H3.3 cells. Conversely, *CDKN1B* encoding P27 was upregulated (Fig4D). The accumulation of P27 was confirmed by western blot analysis and appeared proportional to the percentage of cells in G1 (Fig4E). P27 is known to prevent the G1/S transition by repressing the CDK4-Cyclin D1 (*CCND1*) and CDK2-cyclin E1 (*CCNE1*) protein complexes. S phase entry is mainly under the control of cyclin D/CDK4/6 activation in early G1 leading to activation of CDK2/Cyclin E in late G1 ^26^. Moreover, we also observed at transcriptional level an inhibition of *CCND1* in all DMGs which will also tend to prevent S phase entry (Fig4D).

P27 controls G1 phase along with ERK protein kinase and as mentioned previously, we measured a significant decrease of phospho-ERK in shVRK3 cells (Fig3B).

The modulation of the CDK inhibitor *CDKN1A*, which encodes P21 and is in particular involved in the inhibition of CDK2-cyclin E1 complexes and in G1 block, is more heterogenous between GSC cells. Indeed, *CDKN1A* and *CCNE1* showed an opposite behavior at RNA level following *VRK3* KD: *CDKN1A* was up-regulated in H3.1-mutated cells and slightly down-regulated in H3.3; *CCNE1* was downregulated in H3.1 and slightly upregulated in H3.3 for (Fig4D). WB analysis confirmed the induction of P21 in H3.1-mutated GSC4, and its reduction in GSC1 H3.3-mutated cells, however it also showed to a lesser extent the increase in P21 content in GSC2 (Fig4E).

The decrease in P21 protein content observed specifically in GSC1 cells could result from a smaller decrease of *VRK3* in these cells after transduction and explain that the proportion of cells in S phase in GSC1 cells with shVRK3 remain higher than the other cells (Fig4B).

While a direct binding to CCNB1 and its phosphorylation by VRK3 was previously reported ^16^, we observed here a decrease of *CCNB1* in DMG cells after *VRK3* KD.

CDKs are regulated by several kinases in particular *WEE1,* likewise decreased in shVRK3 cells (FC 0.28; padj 3.07e-15). Additional proteins linked to progression through G1 phase of the cell cycle are greatly modulated following *VRK3* KD (Table S3). Among them, we can mention the strong downregulation of *ANAPC13*, a component of the Anaphase Promoting Complex (FC 0.14; padj 1.57 e-99) that additionally controls progression through mitosis (Fig2D).

### Gene overexpression after *VRK3* KD mainly restricted to metabolism associated genes

As mentioned previously, RNAseq data suggest metabolic reprogramming in cells following *VRK3* repression, as reflected by a significant overexpression of genes involved in ATP-synthesis coupled electron transport and oxidative phosphorylation (OXPHOS) (Fig1D and Fig5A&B). Additionally, *ATP5G3* a metabolism-associated gene is among the top 10 modulated genes (FC 0.1, padj 1.57 e-99, FigS1D & Fig2D). We also noticed a global increase of mitochondrial RNA content in shVRK3 *versus* shCTRL cells (FigS5A). Consequently, we wondered if the overexpression of OXPHOS-related genes resulted from transcriptional modulations only or could also reflect a variation in term of mitochondrial DNA (mtDNA) content as large variations in mitochondria copy number were reported in some tumors ^27^. Additionally, Shen *and coll.* recently reported a decrease of mtDNA copy number in DIPG post-mortem samples *versus* normal brain, but using autopsy samples of frontal lobes consisting primarily of gray matter as reference samples ^28^. We evaluated the mtDNA copy number in DMG-K27 altered primary tumors (n=15) compared to non-tumoral pontine tissue samples (n=6). Despite variability among patients, we measured a tendency for decreased mtDNA content in DIPG either by QPCR (FigS5B) and WES analyses (*data not shown*), confirming the observation of Shen *and coll*. This observation is in accordance with a higher mitochondrial RNA content in non-tumoral pontine tissue samples in comparison with DMG-K27M primary tumors (FigS5C, pvalue 0.00015 Wilcoxon test). However, no modulation of mtDNA content that could explain upregulation of OXPHOS genes was identified by QPCR after *VRK3* repression even 120h after transduction (*data not shown*).

**Fig5.**
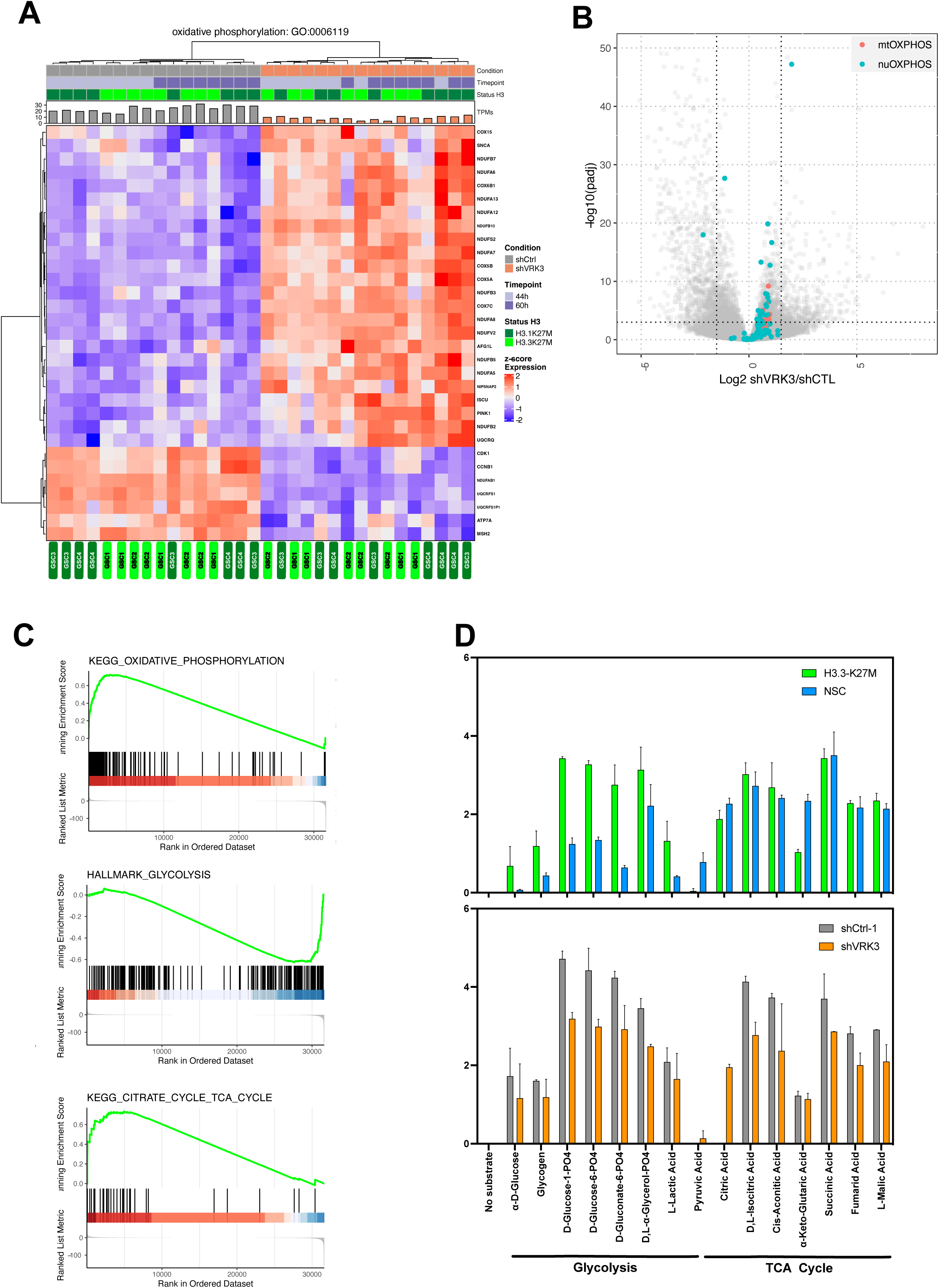
Impact of *VRK3* on metabolism in DMG H3K27-altered. A-Hierarchical clustering and heatmap of modulated genes after *VRK3* KD involved in oxidative phosphorylation (Gene Ontology GO:0006119). B-GSEA plots showing KEGG oxidative phosphorylation, citrate cycle and Hallmark glycolysis pathways in GSC cells following *VRK3* KD. C-Modulation of both nuclear and mitochondrial genes related to OXPHOS. Volcano plot of gene expression in shVRK3 *versus* shCTRL cells plotting antilog of adjusted pvalue on y-axis *versus* log2 Fold Change on x-axis. Nuclear (nuOXPHOS) and mitochondrial (mtOXPHOS) genes are depicted in blue and red respectively. D-The rate of oxidation of a panel of different types of substrates was assessed using Biolog Mitoplate S1 assay. The relative intensity of each substrate is shown as histogram, error bar reflecting the standard deviation of replicate experiments, *i.e.* H3.3-K27M cells (n=2) and NSC cells (n=2), (GSC n=2 for each condition NT, shCTRL1, shVRK3).

Around 10% of OXPHOS genes (13/132 in KEGG oxidative phosphorylation pathway) are encoded by the mitochondrial genome. Looking independently at the modulation of the two subsets of OXPHOS related genes, we identified an upregulation restricted to mtOXPHOS genes in DMG *versus* normal NSC (FigS5D). In contrast, both subsets were significantly upregulated in GSC after *VRK3* depletion (Fig5C) suggesting a specific and global impact of *VRK3* depletion in the induction of this pathway. Among these DEGs, *COX7C* encoding the Cytochrome C oxidase subunits VIIC of the electron transfer respiratory chain (ETC) was one of the most significantly upregulated genes in shVRK3 cells (FC 3.95, padj 6.64E-48)(Fig2D). We then tried to investigate if the deregulation of oxidative phosphorylation following *VRK3* repression was more extensively linked to modifications of cellular metabolism. GSEA analysis of RNAseq data confirmed the enrichment in shVRK3 cells of the ‘oxidative phosphorylation’ geneset (enrichment score (ES) 0.72, qvalue 0.007), highlighted the enrichment of ‘TCA cycle related genes’ (ES 0.73, qvalue 0.35) and a tendency for a depletion of genes involved in ‘glycolysis’ (ES 0.62, qvalue 0.08; Fig5B and FigS5E). Looking at the modulation of individual selected genes in metabolic pathways showed a significant upregulation of majority of genes in OXPHOS, *GALT* in galactose metabolism and *SLC2A3* and *PGAM2* in glycolysis (FigS5E). Other glycolysis-related genes harbored opposite modulation, as they were significantly downregulated in shVRK3 cells: *PGK1*, *PFKFB3* and *PFKFB4*. Of note, PFKFB4 is a regulatory enzyme that synthesizes a potent stimulator of glycolysis and its repression was shown to suppress tumor growth and metastasis in breast cancer^29^, and is considered as a key molecule for survival of cancer stem cells in glioblastoma^30^. Indeed, PFKFB4 enhances G1/S transition by increasing CDK6 level^31^. In glioma, E2F2-dependent overexpression of *PFKFB4* leads to PI3K/AKT pathway activation and its repression was shown to impair cell growth ^32^(*E2F2* FC 0.117, padj 1.11 e-08; *PFKFB4* FC 0.29, padj 4.5 e-08).

We next investigated mitochondrial metabolic activity and substrate preference including TCA cycle intermediates, hexoses, trioses, amino acids and fatty acids. The rate of substrate oxidation of amino acid and fatty acid substrates were close to background level, and appeared significantly decreased for glycolysis and TCA cycle substrates after *VRK3* KD (Fig5D). These results confirmed a lower glycolysis in GSC after *VRK3* KD, as suggested by GSEA analysis. Interestingly, we observed a global increase of glycolysis in GSC *versus* NSC, *VRK3* repression tending to reset this metabolic pathway. The decrease in the rate of oxidation of several substrates of the TCA cycle, such as fumaric and isocitric acids suggests a reduction of this metabolic activity despite an enrichment of several upregulated genes in shVRK3 cells highlighted by GSEA.

The imipridone ONC201, a potent activator of the mitochondrial Clp protease proteolytic subunit (ClpP), was shown to inhibit cell proliferation and induce cell death in many cancers including GBM ^33^ and one DMG patient ^34^. ONC201 lead to a decrease expression of respiratory chain proteins and subsequently impair OXPHOS that results in mitochondrial dysfunction and tumor cell death.

We tested ONC201 in our GSC models (n=10, 5 H3.1- and 5 H3.3-mutated cells) and found IC50 ranging from 0.95 to 1.7 µM (median 1.35 µM, *data not shown*). This significant efficiency appeared independent of *EGFR* expression level in the corresponding primary tumors despite the fact that its efficiency was reported to be inversely proportional to *EGFR* expression in GBM ^35^.

## Discussion

The gene expression study after *VRK3* KD gives some clues to explain the profound impact of VRK3 deprivation on DMG cell fitness. There are only few differences between the two main subgroups of DMG-H3K27 altered, *i.e.* H3.1- and H3.3-K27M DMG, despite that the H3.3 GSC models used are *TP53*-mutated and that VRK3 was shown to interact with TP53 protein. The massive gene downregulation observed after *VRK3* inhibition affect mainly genes involved in chromatin segregation. Chromatin condensation, one of the crucial steps in mitosis, is known to be enhanced via phosphorylation of histone H3 on Ser10. In accordance, we observed a significant decrease of H3S10P-positive cells in shVRK3 cells as well as modulation of several key regulators of H3S10P (AURKA, B, MSK2, 14-3-3). Phosphorylation of H3S10 was shown to be critical during neoplastic transformation, and the steady state level of H3S10P is elevated in oncogene-transformed cells and human tumor cell lines ^36, 37^. Accordingly, we observed a higher content of H3S10-positive cells in H3.3-K27M GSC *versus* NSC.

VRK3 protein appears to be mostly located in the nucleus, with the majority unbound to chromatin in DMG-K27M. Consequently, we can suppose that VRK3 is not directly involved in H3S10 phosphorylation. In contrast, a significant proportion of the nuclear protein VRK1 is bound to chromatin in DMG, as reported in fibroblasts ^17^ which progressively regulate Histone H3 during chromatin compaction in G2/M and mitosis ^38^. First, VRK1 phosphorylates H3 on Thr3 allowing its binding to the survivin/BIRC5 complex and the recruitment of AURKB, which then lead to H3S10 phosphorylation ^8, 10^. The reduction in H3S10 phosphorylation following *VRK3* KD could at least partially results from a decrease of *VRK1*.

In mammals, H3 phosphorylation occurs at two serine residues, S10 and S28, which can be mediated by histone kinases including MSK1 and AURKB. In shVRK3 cells, both H3 phosphorylations are decreased in parallel with *AURKB* repression.

H3S10P is also involved in chromatin dynamics and was reported to prevent deposition of H3K9me2 ^22^. VRK3 is known to interact directly with several histone proteins and chromatin modifiers (Lee 2017). Here, we have shown modulation of genes involved in histone acetylation 60h after *VRK3* KD as well as an inhibition of a component of PRC2 complex (JARID2), and a strong decrease of *SMARCA1* encoding an ATPase of the ISWI subfamily of chromatin remodelers SWI/SNF (FC 0.06, padj 5.07 e-78). All these observations support its role in chromatin dynamics beyond the modulation of H3S10P.

Despite the large effect at transcriptional level on genes involved in chromosome segregation, the major phenotypical impact of *VRK3* depletion is a G1 block. Indeed, many genes linked to cell proliferation are downregulated, explaining the G1/S transition arrest observed (*i.e.* a significant increase of cells in G1 phase and loss of cells in S phase). Another study reported the inhibition of cell growth following *VRK3* silencing ^16^. However, it was associated with S phase and G2/M phase arrests in liver cancer. This discrepancy could reflect a substantially distinct role of VRK3 depending on the cellular context, even though we also observed minor modulations of genes involved in G2/M transition. These authors have also shown that VRK1 plays a crucial role in G1/S transition and that its depletion induces a G1 arrest by downregulating cyclin D1 and phospho-RB, and upregulating P21 and P27 in hepatocellular carcinoma ^16^. As we have shown that *VRK3* KD leads to a substantial decrease of VRK1 associated with the upregulation of P21 and P27 and downregulation of *CCND1*, one can hypothesize that the observed blockade of DMG cells cycle upon *VRK3* KD goes through VRK1 downregulation.

Recently VRK1 was shown as a synthetic lethal target in VRK2-deficient glioblastoma ^18^ and more globally in VRK2-promoter methylated cancers of nervous system ^39^, including DMG. In the second study, they showed that VRK1 dependency was inversely correlated with expression of *VRK2*, and that *VRK2* promoter methylation was associated with low *VRK2* expression level mainly in G34R pediatric glioma. However, *VRK1* KO led to cell death without alteration of cell cycle profile in DMG models. In our models we did not observe an hypermethylation of *VRK2* promoter (figS3M) and observed a higher correlation between *VRK3* and *VRK1* expression in comparison to *VRK3* and *VRK2*. Our previous work has showed that our models of K27M-DMG are more dependent on *VRK3* than *VRK1* or *VRK2* ^7^. Indeed, we also observed that *VRK1* extinction by RNA interference affect DMG survival but to a significant lesser extent than VRK3. This observation reflects that it is unlikely that VRK1 can fully compensate VRK3 function, making of VRK3 a more interesting target in DMG-K27M. It would be interesting to investigate if *VRK3* depletion is also synthetic lethal in the other pediatric tumor entities without hypermethylation of VRK2 promoter, and in particular posterior fossa type A ependymoma and infant high-grade glioma (figS3M). VRK1 and VRK3 share some common substrates ^16^, but beyond their overlapping role on cell cycle progression, VRK3 presents other specific function impairing DMG-K27M survival. Interestingly, the comparison of VRK1 and VRK3 interactomes uncovers that VRK3 interacts specifically with proteins located in endoplasmic reticulum and mitochondria ^16^.

Metabolic reprogramming is a hallmark of cancer cells, that can result in many tumors from a variation in mtDNA copy number content. A decrease of mtDNA copy number is observed in the majority of tumors ^27^. Similarly, DMG harbor a lower mtDNA copy number than control NSC cells which is not altered by *VRK3* repression.

Most tumors, including adult glioblastoma (GBM), showed important indicators of reduced OXPHOS system activity compared with that of the electron transport system. Indeed, glioma cells predominantly rely on glycolysis instead of oxidative phosphorylation to regenerate ATP ^40–42^, a phenomenon named the Warburg effect ^43^, resulting from the abnormal availability of respiratory substrates. Accordingly, in our DMG-H3K27M GSC models we observed a disruption of energy metabolism favoring glycolysis over OXPHOS in comparison to our NSC control cells. We partially validated the results of Chung and coll., generated by comparing isogenic NSC cells transduced or not with H3K27M construct, with only an increase of glycolysis in K27M cells ^44^. Knocking-down *VRK3* tends to revert this metabolic phenotype and leads to growth arrest. *VRK3* KD affects metabolism in particular by upregulation of the majority of respiratory electron transport chain as well as OXPHOS related genes. Shen *and coll.* have shown that PDK inhibitors are also able to revert the Warburg effect by stimulating OXPHOS in DMG and restore mitochondrial metabolism, which was associated with a decrease cell viability ^28^. HDAC inhibitors revert the Warburg effect through a reduction in glycolysis leading to energy deprivation, which in turn leads to enhancement of oxidative metabolism ^45^. Additionally, it was recently shown in adult GBM that the activation of ClpP through utilization of the second-generation compounds (ONC206 and ONC212) in combination with pharmacological inhibition of HDAC1/2 cause synergic reduction of GBM viability ^46^. Lately, ONC201, a potent activator of the mitochondrial ClpP leading to OXPHOS impairment and decrease in enzymatic activity of respiratory chain complexes I, II, and IV, was proposed as an anticancer strategy in K27M-DMG ^47^. Accordingly, ONC201 appeared efficient *in vitro* in all DMG-K27M GSC models tested. This family of compound was however less efficient in GBM unless OXPHOS was upregulated ^46^, and ONC201 has been shown to be more effective in AML which were dependent to OXPHOS ^48^. Our data support that DMG K27M present a dependency on OXPHOS as well and that strong up-regulation of OXPHOS genes is associated with impairment of their growth after *VRK3* KD. It could thus be anticipated that combining ONC201 with pharmacological targeting of VRK3 would be synergistic.

## Materials and Methods

### Cell and culture

Human Glioma Stem Cell primary culture derived in the lab corresponding to three H3.3-K27M (GSC1, GSC2, GSC5) and 2 H3.1-K27M (GSC3, GSC4) cellular models were grown in Neurocult^TM^ medium supplemented with heparin (2 μg/mL, Stemcell Technologies), EGF/FGF (20ng/mL, Miltenyi) at 37°C and 5% CO2 using laminin-coating. Normal stem cell (NSC3, NSC4, NSC5) derived as previously published ^7^ were grown in the same culture conditions. For GSC cells, PDGF (10 ng/mL, Miltenyi) was added to the culture medium. HEK293T and HCT116 were cultivated in DMEM supplemented with 10% fetal bovine serum and 1% Penicillin/Streptomycin (Life technologies) at 37°C and 5% CO2. For all cell types, medium was renewed in a two-days basis. All the samples were obtained with the written informed consents for the biopsy of subjects according to Helsinki declaration.

### Cell cycle analysis by flow cytometry

Cells were exposed during 2 hours to 10 μmol/L of EdU supplied with Click-iT Flow Cytometry Assay Kit (ThermoScientific) and then detached using accutase® (ThermoFischer Sicentific). EdU was incorporated into cellular DNA during replication to tag cells in S phase. Total DNA was stained with FxCycleTM Violet stain (Thermofisher Scientific). Samples were acquired using an LSR Fortessa flow cytometer (BD Biosciences), and analysis was performed with Flowjow software (Flowjow 10.8.1).

### Lentiviral production and transduction

Lentiviral particles were produced in HEK293T cells using psPax2 and pMD2.g second-generation packaging plasmids (Addgene #12260, #12259) with jetPRIME Polyplus transfection reagent, and lentiviral titers were then determined by fluorescence assay. GSCs were transduced at a multiplicity of infection (M.O.I.) of 3 with shVRK3-1, shVRK3-4, shCtrl-1 (negative control vector containing a nonhairpin insert Addgene #1864) and shCtrl-2 (MISSION® pLKO.1-puro non-mammalian shRNA Control Plasmid DNA, SHC002, Sigma-Aldrich).

### Gene expression of tumor cells after VRK3 silencing with RNA-Sequencing

Five hundred thousand cells (GSC1, GSC2, GSC3, GSC4) were transduced ^7^. Total RNA was extracted with Direct-zol RNA micro prep (Zymo research) at 44 hours and 60 hours post-transduction.

The RNA integrity (RNA Integrity Score ≥ 7.0) was checked on the Agilent 2100 Bioanalyzer and quantity was determined using Qubit (Invitrogen). SureSelect Automated Strand Specific RNA Library Preparation Kit was used according to manufacturer’s instructions with the Bravo Platform. Briefly, 50 to 200 ng of total RNA per sample was used for poly-A mRNA selection using oligo(dT) beads and subjected to thermal mRNA fragmentation. The fragmented mRNA samples were subjected to cDNA synthesis and were further converted into double stranded DNA using the reagents supplied in the kit, and the resulting dsDNA was used for library preparation. The final libraries were bar-coded, purified, pooled together in equal concentrations and subjected to paired-end sequencing on Novaseq-6000 sequencer (Illumina). Raw data quality checking was performed with FastQC and pseudo mapping with Salmon (v0.9.1) using Gencode annotation (v27) based on hg38 reference genome (GRCh38); Genes with less than 10 reads in the resulting raw count matrix were filtered. After variance-stabilizing transformation and removal of sample model bias with limma removeBatchEffect, DESeq2 was used with default parameters to select differentially expressed genes (DEGs) with a pvalue threshold of 0.001 after Benjamini and Hochberg correction. Sample hierarchical clustering was performed using complexHeatmap function with Euclidian distance and complete linkage as aggregation method. For functional annotation and enrichment analysis, we used GeneTonic package, ClusterProfiler package (GO from org.Hs.eg.db GOSOURCEDATA = 2021-02-01) and Gene Set Enrichment Analysis (GSEA) to identify gene sets from the MSigDB v7.4 database (C5 (GO:BP), C2 (KEGG) and HALLMARK catalogue) overrepresented among log2FC ranked genelists, using Benjamini and Hochberg procedure as multiple testing correction. Enrichmap was obtained by computing Jaccard distance between top 100 GOenrich genesets. The communities in Enrichmap were identified by Louvain clustering. Hierarchical clustering of selected gene subsets was conducted using z-score, Pearson correlation and ward D2 as agglomeration criteria.

### DNA Methylation Array Processing

DNA methylation data of the samples of Capper et al ^49^ and Castel et al ^50^ analysed were performed in R V4.0.3. Raw signal intensities were obtained from IDAT files using the minfi Bioconductor package version 1.36.0. Each sample was individually normalized by performing a background correction and a dye-bias correction. Probes located on sex chromosomes, not uniquely mapped to the human reference genome (hg19), containing single nucleotide polymorphisms and that are not present in both EPIC and 450k methylation array were removed. Subsequently, a batch effect correction for the type of material tissue (FFPE/frozen) was performed using limma package v3.30.11). The remaining probes were sorted by standard deviation, the 10,000 most variable were selected and used to compute the 1-variance weighted Pearson correlation between samples. The distance matrix was used as input for t-distributed stochastic neighbor embedding (t-SNE) from Rtsne package, with the following non-default parameters: theta = 0, pca = F, max_iter = 2500 and perplexity = 20. The beta value of the probe cg26093711, localized upstream of the *VRK2* gene, where generated with the function getBeta() from the minfi package

### Phospho-Array

Two-hundreds microliter of lysis buffer are added to cells after VRK3 silencing and two-and-a-half million cells were scrapped (99003, TPP). Proteins were then extracted and purified using the kit Antibody Array Assay Kit (KAS02) according to manufacturer recommendations (Full moon BioSystem). Fifty μg of proteins were biotinylated, deposited onto the Phospho Explorer Antibody Array containing 1,318 antibodies, hybridized and washed according to manufacturer recommendations. The slides were scanned on a G2505C Agilent scanner at an excitation wavelength of 550nm at a resolution of 10 μm, and images analyzed with Agilent Feature Extraction 12.0 (Agilent Technologies, Inc 2015). The fluorescence signal of each antibody (‘processed signal’) was corrected by subtracting of the local background. Fold change of corrected signal intensity were analyzed in shVRK3-4-transduced cells *versus* corrected signal intensity in shCtrl2 -transduced cells.

### Protein extraction and immunoblotting

For classical protein extraction, cells were lysed as described previously ^7^ in RIPA lysis buffer. For acidic protein extraction, we used Histone Extraction Kit (Abcam, cat # ab113476) and followed manufacturer protocol. Protein quantification was performed by spectrophotometry with PierceTM BCA Protein Assay Kit (ThermoFischer Scientific, cat # 23225) and proteins were separated on a 4% to 20% polyacrylamide gel (Biorad), and transferred to PVDF membrane (Bio-rad) with a Trans-Blot Turbo system (Bio-Rad). Membranes were incubated overnight at 4°C with primary antibody (VRK3 #HPA056489 Sigma, B-actine #51255 Cell signaling, Lamin # ab16048 Abcam, H3S10P #05-806 Millipore, H4 #07-108 Millipore, H3S28P #07-145 Millipore, VRK1 #HPA000660 Sigma, MSK2 #3679S Cell signaling), and 1 hour at 20°C with horseradish peroxidase-linked secondary antibody for (# 7074S (1/2000), #7076S (1/5000) Cell Signaling Technology), #0500-0099 (1/50000, Biorad). Proteins were enhanced by chemiluminescence reagent (Biorad) and analyzed with ChemiDocTM XRS+ System (Biorad) with Image Lab 4.1 Software (Bio-Rad).

### Immunofluorescence

For immunofluorescence, cells were plated the day before fixation on coverslip coated with 0,1 mg/mL Matrigel (Corning). Fixation was performed 5 minutes with PBS1X/4%paraformaldehyde, permeabilization for 5 minutes with 0,5% Triton X-100, and blocking for 30 minutes in 1X PBS/5% normal goat serum (ThermoFisher). Cells were incubated overnight at room temperature (RT) with anti-VRK3 (1/1000 #HPA056489, Sigma) and anti-Phalloidine (1/200 #8940S, Cell signaling) and 45 minutes with secondary antibodies (Alexa fluor, Invitrogen 1:800) and Hoechst 33342 (5 ug/mL #H3570, ThermoFisher) before washes. Slides were mounted with fluoromount-G (SouthernBiotech). Images were acquired with Leica SP8 confocal microscope. For immunofluorescence in suspension, cells were plated the day before transduction and fixed 80 hours post-transduction according to thermofisher’s protocol. Cells were incubated with antibodies anti-VRK3 (#HPA056489, Sigma), anti-H3S10P (1/1000 #05-806, Millipore), anti-H3S28P (1/1000 #07-145 Millipore) overnight at 4°C and with secondary antibodies (Alexa fluor, Invitrogen 1:800) during 1h at RT. Samples were acquired using an LSR Fortessa flow cytometer (BD Biosciences), and analysis was performed with Flowjow software (Flowjow 10.8.1).

### RT-qPCR

Total RNAs were extracted and purified with Direct-zol^TM^ RNA MicroPrep (Zymo Research) according to manufacturer’s instructions under RNase free conditions. Five hundred ng of total RNA was reverse-transcribed using Revertaid first strand cDNA synthesis kit (Thermo Fisher Scientific). Real time quantification was performed in triplicate with ViiA Real-Time PCR system (ThermoFisher Scientific). Each sample was analyzed in triplicates. Expression of *TBP* (TATA-binding protein) was used as an internal loading control and 2**^-ΔΔ^**^CT^ method was used for relative expression computation. Primer sequences are summarized in Table S1.

### Mitotracker staining

Cells were detached with accutase and washed two times with PBS 1X. Cells were then exposed to Green mitotracker^TM^ (Thermofisher, #M7414) during 30 minutes at 37°C and washed two times with PBS 1X. Samples were acquired using an LSR Fortessa flow cytometer (BD Biosciences), and analysis was performed with Flowjow software.

### Mitochondrial function assays

Mitochondrial function assays were performed using MitoPlate S-1(Biolog Inc., Hayward, CA). The substrates on MitoPlate S-1 were first dissolved by incubating the plate with 30 µl of Assay Mix, consisting of 2x BMAS, 6x Redox Dye MC and saponin (30 µg/ml) necessary for cell permeabilization in a 5% CO2 incubator at 37°C for 1 h before inoculating 400,000 cells per well in a volume of 30 µl. Color changes are read kinetically during 24h and are performed with OmniLog instrument (Biolog Inc., Hayward, CA). Signal intensity was measured at 24h and corrected by background subtraction with the negative control condition without substrate. The relative intensity was computed by dividing each intensity by the global mean of a particular sample.

### Drug evaluation

GSC and NSC cells were plated in duplicates at 10.000 - 15.000 cells/cm2 in 96-well plates in 100 μl of complete GSC medium. Twenty-four hours after seeding, Ro31220 (Calbiochem, cat # 557520), SB747651A (Tocris, cat # 4630) or vehicule DMSO (Sigma-Aldrich) were added using the D300E digital dispenser (TECAN). ONC201 (TIC10, Selleckchem cat #S7963) was added with a range of 0.25-8µM. Cell proliferation was assessed during 7 days after treatment by videomicroscopy with an Incucyte Zoom or S3 (Sartorius). Cell confluence was determined by CellPlayer Analysis software (Essen Bioscience) and normalized against the initial time point. Inhibitory Concentration (IC50) values were obtained by plotting logarithmic concentrations of the drug against the area under the curve.

## Figure legends

**FigS1**

A-*VRK3* expression level in RNAseq (tpm) is significantly decrease in shVRK3 samples (orange) in comparison to shCTRL samples (dark gray, Mann-Whitney, pvalue 0.0159) both for H3.1-(light green) and/or H3.3-mutated cells (dark green). NT, non-transduced cells in light grey

B. Distribution of Differentially Expressed Genes (DEGs) following *VRK3* KD selected with an adjusted pvalue threshold of 0.001 according to the time post-transduction.

C. Distribution of DEG following *VRK3* KD selected with an adjusted pvalue threshold of 0.001 according to H3.1 and H3.3 mutational status (*i.e.* dark green for DEGs found specifically in H3.1-K27M cells, light green for DEGs found specifically in H3.3-K27M cells and red for common DEGs).

D. Volcano plot of gene expression in shVRK3 cells 60h *versus* shVRK3 cells 44h post-transduction plotting antilog of adjusted pvalue on y-axis *versus* log2 Fold Change (FC) on x-axis. DEGs associated with adjusted pvalue <0.001 are color-coded in red.

E. Volcano plot of gene expression in shVRK3 cells *versus* shCTRL cells, plotting antilog of adjusted pvalue on y-axis versus log2 Fold Change on x-axis. DEGs associated with adjusted pvalue <0.001 are color-coded in red.

**FigS2**

Representation of the four distinct communities identified by Louvain clustering among the enrichment map presented in figure 2.

The size of the node encodes the information of the number of DEGs while the color is representative of the computed Z score for each set. The color is indicative of the significance of expression changes of genes assigned to a particular set.

**FigS3**

A- Modulation of phospho-MSK2 level depicted by phospho-array analysis 72h after transduction with shCTRL-2 or shVRK3-4 in H3.3-K27M GSC2 (ligh green) and H3.1-K27M (dark green) GSC4 cells. Error bars represent standard deviation of experimental duplicates.

B- Dose-response results displaying the effect of MSK2 inhibitor on NSC and GSC proliferation. Proliferation was monitored by video microscopy for 168 hours. Normalized areas under the curve (AUC) were plotted against logarithmic concentrations of Ro318220 drugs to determine the IC50. H3.3-K27M GSCs are shown in light green and H3.1-K27M GSC in dark green.

C- Mean of percentage of H3S10P-positive cells assessed by immunofluorescence in several experimental replicates (*i.e.* GSC1/2/4 n=5, GSC3 n=2 and for control samples NSC4/5 n=4,). Error bars correspond to standard deviation of experimental replicates. * t-test pvalue<0.05, ** t-test pvalue<0.005.

D- Western Blot analysis showing levels of H3S10P in GSC1, GSC2 and GSC5 (H3.3-K27 model, light green) and NSC3 with or without nocodazole treatment. Histone H4 was used as loading control.

E- Modulation of the percentage of H3S10P-positive cells after *VRK3* KD in several experimental replicates (*i.e.* GSC1 n=3, GSC2/3 n=1, GSC4 n=2). * t-test pvalue<0.05, ** t-test pvalue<0.005, *** t-test pvalue<0.0005.

F- Mean of percentage of H3S28P-positive cells assessed by immunofluorescence in several experimental replicates (*i.e.* GSC1/2/4 n=4, GSC3 n=1 and for control samples NSC4/5 n=3). Error bars correspond to standard deviation of experimental replicates. * t-test pvalue<0.05, ** t-test pvalue<0.005,

G-Modulation of the percentage of H3S28P-positive cells after *VRK3* KD in several experimental replicates (*i.e.* GSC1 n=3, GSC2/3 n=1, GSC4 n=2). * t-test pvalue<0.05, ** t-test pvalue<0.005, *** t-test pvalue<0.0005.

H- Analysis of biological process enrichment of genes encoding VRK3 interacting proteins from Lee and coll. (Lee et al. 2017) using enrichGO function of cluster profiler package. Adjusted pvalue from Fisher exact test are color-coded.

I- Hierarchical clustering of gene encoding VRK3 interacting proteins identified by Lee and coll (Lee et al. 2017). Heatmap show RNAseq expression level (Z-score on normalize expression matrix) and on the left of the heatmap differentially expressed genes identified using either an adjusted pvalue threshold of 0.01 or 0.001 are indicated as red squares.

J-K- Analysis of the subcellular location of VRK1 and VRK3 by Western Blot in total, cytosolic, nuclear soluble and nuclear insoluble fractions in GSC2 and GSC4 cells. β–Actin, lamin B1 and H4 were used as loading control for total/cytosol, nuclear soluble and nuclear insoluble fractions respectively.

L*-* Dot plot presenting *VRK1 vs. VRK3* expression levels on left panel and *VRK2 vs. VRK3* expression levels on the right panel from RNAseq data of primary DMG (tpm). Spearman (R) and Pearson correlation (p) are indicated.

M- t-SNE analysis of DNA methylation profiles of 974 gliomas selected reference samples. Left panel shows the distribution of the samples colored by their methylation beta value for the probe cg26093711 located in *VRK2* promoter region. Right panel shows the same samples colored by tumor entity.

**FigS4**

Modulation of *VRK3* expression during cell cycle assessed by qRT-PCR (A) or immunofluorescence (B). The mean fluorescence intensity of cells across cell cycle was corrected by fluorescence intensity in secondary antibody alone. One-way ANOVA was performed only for GSC cells with at least two independent experiments. One-way ANOVA ** pvalue < 0,01 **** pvalue < 0,0001. Light green, H3.3-K27M GSCs and dark green H3.1-K27M GSC.

**FigS5**

A- Boxplot reflecting global level of expression of mitochondrial RNA in GSC transduced with shCTRL or shVRK3 (tpm from RNAseq data) (Wilcoxon test *pvalue<0.05, **<0.005).

B- Relative mtDNA copy number measured by qPCR in primary tumors (pontine DMG H3K27-altered n=15) and non-tumoral primary pontine tissue samples (NC ou NonContrib, n=6) of DMG-K27M patients. Error bars represent standard deviation between samples.

C- Violin plot of global level of expression of nuclear and mitochondrial transcriptomes in DMG-K27M primary tumors (n=15) and non-tumoral primary pontine tissue samples (NonContrib, n=3) (tpm from RNAseq data). Error bars represent standard deviation between samples.

D- Volcano plot of nuclear and mitochondrial genes related to OXPHOS in GSC cells *versus* NSC cells. Nuclear (nuOXPHOS) and mitochondrial (mtOXPHOS) genes are depicted in blue and red respectively.

E- Barplots presenting the fold change of metabolism genes between normal and glioma stem cells (green) as well as in GSC after *VRK3* KD (orange). Benjamini and Hochberg adjusted pvalue * < 0.05, ** < 0.01, *** < 0.001.

## Acknowledgements

DC, JG, MAD acknowledge financial support from charities Lisa For Ever, l’Etoile de Martin and La marche de l’Ecureuil. We acknowledge the use of the bioresources of the PRB Tumorothèque Necker and Necker Imagine DNA biobank (BB-033-00065). The authors are grateful to the Necker hospital tumor and DNA banks and the Necker operating room nurses/assistants for their technical assistance. TK was supported by the charity “Imagine for Margo” and by the Fondation Gustave Roussy “Guérir Les Cancers des Enfants au XXIe siècle (GLCE)”. This project was supported by grant “ Taxe d’apprentissage Gustave Roussy – 2018 - CE”. We thank N. Droin and the genomic core facility of Gustave Roussy.

