## Supplementary material for "VRK3 depletion induces cell cycle arrest and metabolic reprogramming of Pontine Diffuse Midline Glioma (DMG)-K27 altered cells": FigS1, FigS2, FigS3, FigS4, FigS5

### Figure S1

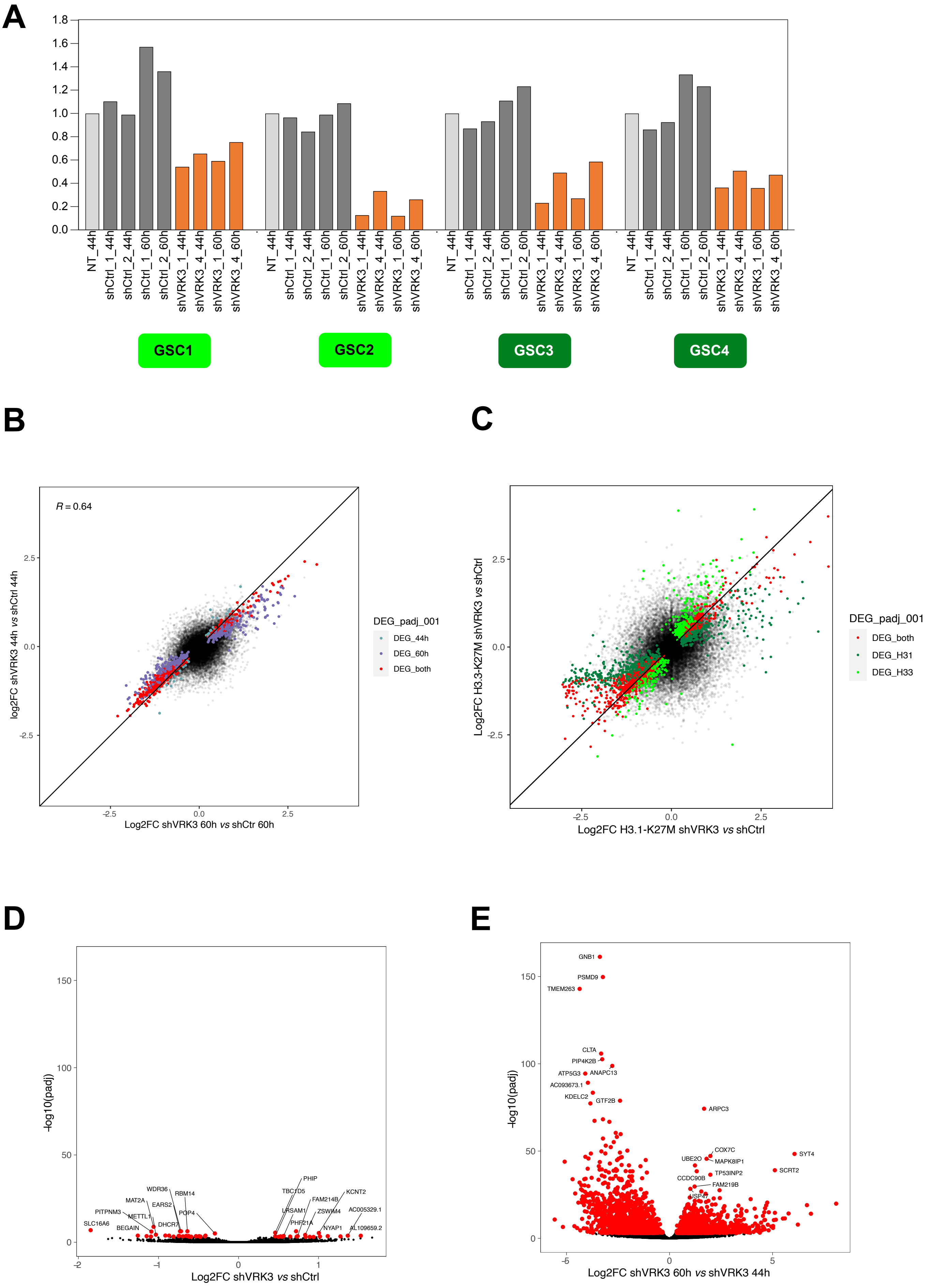

A
NT_44h
shCtrl_1_44h
shCtrl_2_44h
shCtrl_1_60h
shCtrl_2_60h
shVRK3_1_44h
shVRK3_4_44h
shVRK3_1_60h
shVRK3_4_60h
GSC1
GSC2
GSC3
GSC4
B
R = 0.64
log2FC shVRK3 44h vs shCtrl 44h
Log2FC shVRK3 60h vs shCtr 60h
DEG_padj_001
DEG_44h
DEG_60h
DEG_both
C
Log2FC H3.3-K27M shVRK3 vs shCtrl
Log2FC H3.1-K27M shVRK3 vs shCtrl
DEG_padj_001
DEG_both
DEG_H31
DEG_H33
D
-log10(padj)
Log2FC shVRK3 vs shCtrl
SLC16A6
PITPNM3
MAT2A
WDR36
RBM14
EARS2
METTL1
BEGAIN
DHCR7
POP4
PHIP
TBC1D5
FAM214B
KCNT2
LRSAM1
ZSWIM4
AC005329.1
PHF21A
NYAP1
AL109659.2
E
-log10(padj)
Log2FC shVRK3 60h vs shVRK3 44h
GNB1
PSMD9
TMEM263
CLTA
PIP4K2B
ATP5G3
ANAPC13
AC093673.1
KDELC2
GTF2B
ARPC3
COX7C
MAPK8IP1
UBE2O
TP53INP2
CCDC90B
FAM219B
USP47
SYT4
SCRT2

### Figure S2

#### 1- Regulation of cell cycle transition

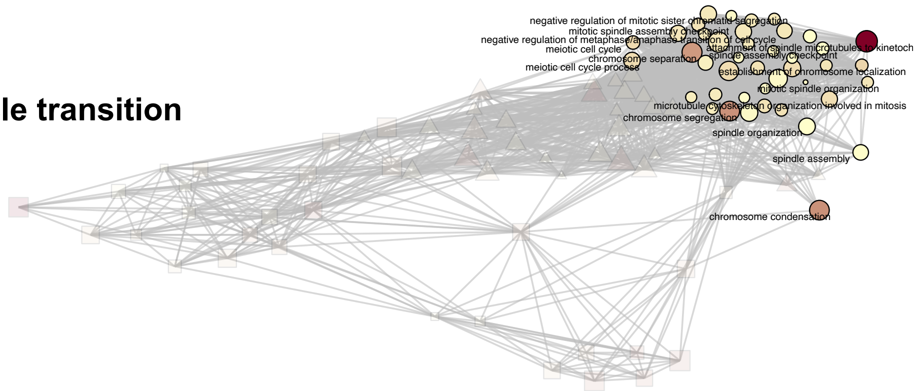


#### 2- Telomere organization

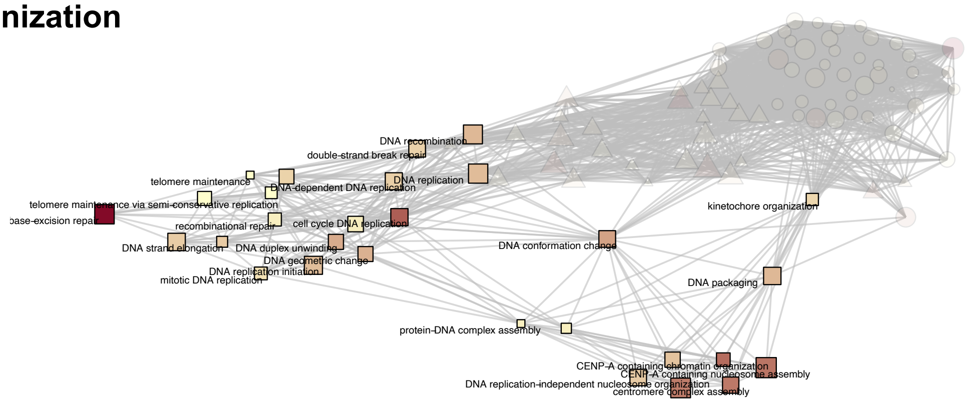


#### 3- Kintochore and centromeric chromatin organization

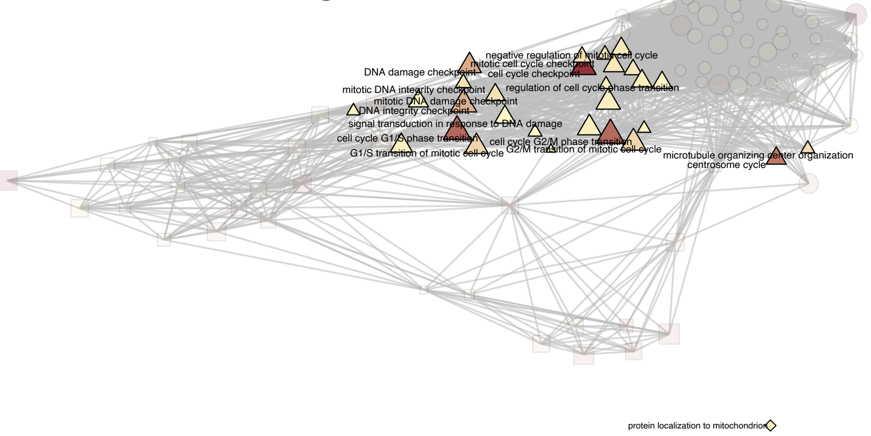


#### 4- Localization to protein mitochondrion

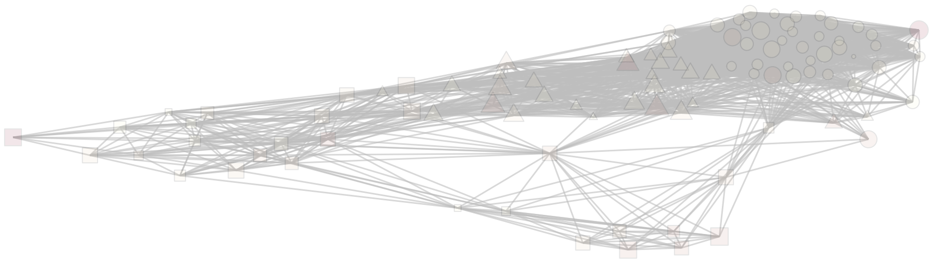

### Figure S3-A

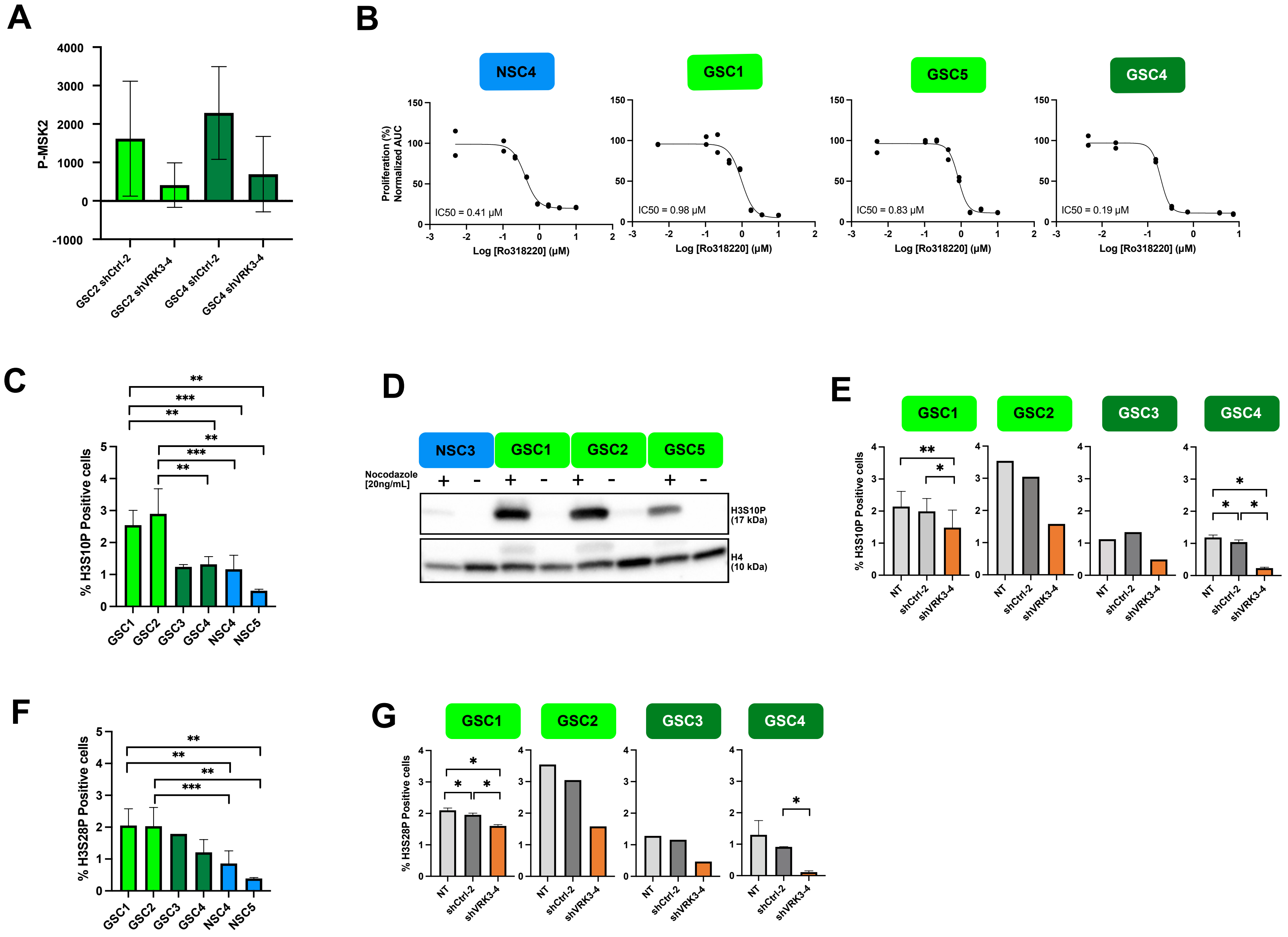

A
P-MSK2
4000
3000
2000
1000
0
-1000
GSC2 shCtrl-2
GSC2 shVRK3-4
GSC4 shCtrl-2
GSC4 shVRK3-4
B
NSC4
GSC1
GSC5
GSC4
Proliferation (%)
Normalized AUC
IC50 = 0.41 µM
IC50 = 0.98 µM
IC50 = 0.83 µM
IC50 = 0.19 µM
Log [Ro318220] (µM)
C
% H3S10P Positive cells
GSC1
GSC2
GSC3
GSC4
NSC4
NSC5
D
NSC3
GSC1
GSC2
GSC5
Nocodazole
[20ng/mL]
H3S10P
(17 kDa)
H4
(10 kDa)
E
GSC1
GSC2
GSC3
GSC4
% H3S10P Positive cells
NT
shCtrl-2
shVRK3-4
F
% H3S28P Positive cells
GSC1
GSC2
GSC3
GSC4
NSC4
NSC5
G
GSC1
GSC2
GSC3
GSC4
% H3S28P Positive cells
NT
shCtrl-2
shVRK3-4

### Figure S3-B

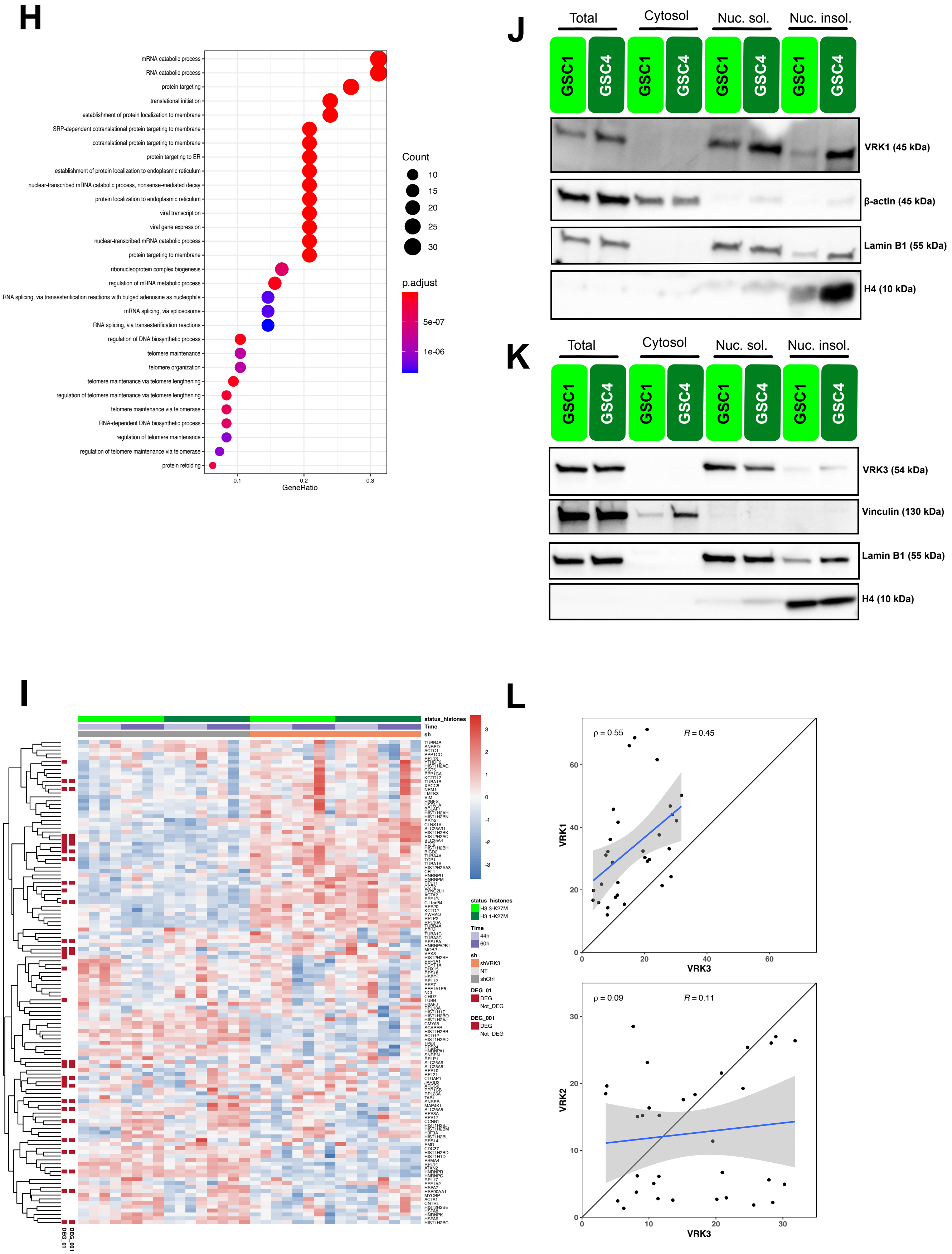

H
mRNA catabolic process
RNA catabolic process
protein targeting
translational initiation
establishment of protein localization to membrane
SRP-dependent cotranslational protein targeting to membrane
cotranslational protein targeting to membrane
protein targeting to ER
establishment of protein localization to endoplasmic reticulum
nuclear-transcribed mRNA catabolic process, nonsense-mediated decay
protein localization to endoplasmic reticulum
viral transcription
viral gene expression
nuclear-transcribed mRNA catabolic process
protein targeting to membrane
ribonucleoprotein complex biogenesis
regulation of mRNA metabolic process
RNA splicing, via transesterification reactions with bulged adenosine as nucleophile
mRNA splicing, via spliceosome
RNA splicing, via transesterification reactions
regulation of DNA biosynthetic process
telomere maintenance
telomere organization
telomere maintenance via telomere lengthening
regulation of telomere maintenance via telomere lengthening
telomere maintenance via telomerase
RNA-dependent DNA biosynthetic process
regulation of telomere maintenance
regulation of telomere maintenance via telomerase
protein refolding
0.1
0.2
0.3
GeneRatio
Count
10
15
20
25
30
p.adjust
5e-07
1e-06
J
Total
Cytosol
Nuc. sol.
Nuc. insol.
GSC1
GSC4
VRK1 (45 kDa)
β-actin (45 kDa)
Lamin B1 (55 kDa)
H4 (10 kDa)
K
Total
Cytosol
Nuc. sol.
Nuc. insol.
GSC1
GSC4
VRK3 (54 kDa)
Vinculin (130 kDa)
Lamin B1 (55 kDa)
H4 (10 kDa)
I
status_histones
Time
sh
status_histones
H3.3-K27M
H3.1-K27M
Time
44h
60h
sh
shVRK3
NT
shCtrl
DEG_01
DEG
Not_DEG
DEG_001
DEG
Not_DEG
DEG_001
DEG_01
L
ρ = 0.55
R = 0.45
VRK1
VRK3
ρ = 0.09
R = 0.11
VRK2
VRK3

### Figure S4

## A

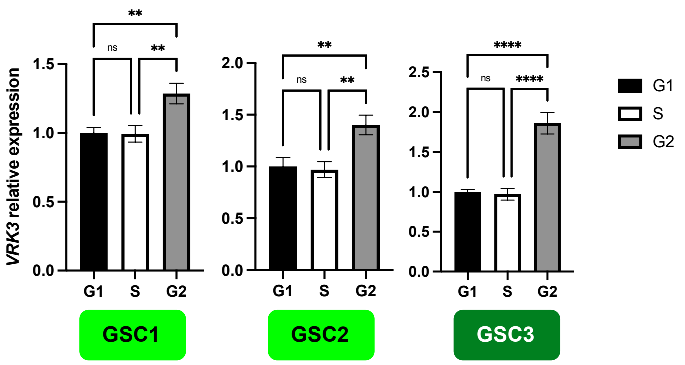

VRK3 relative expression
G1
S
G2
ns
**
****
GSC1
GSC2
GSC3


## B

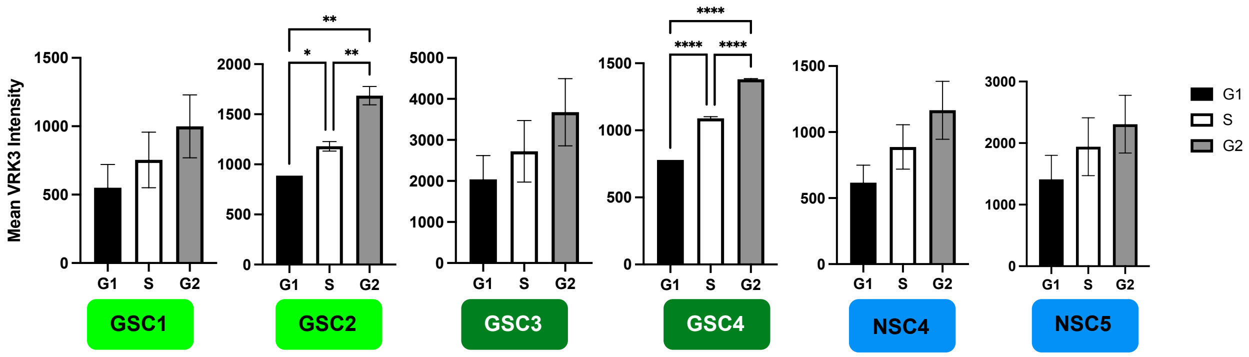

Mean VRK3 Intensity
G1
S
G2
*
**
****
GSC1
GSC2
GSC3
GSC4
NSC4
NSC5

### Figure S5

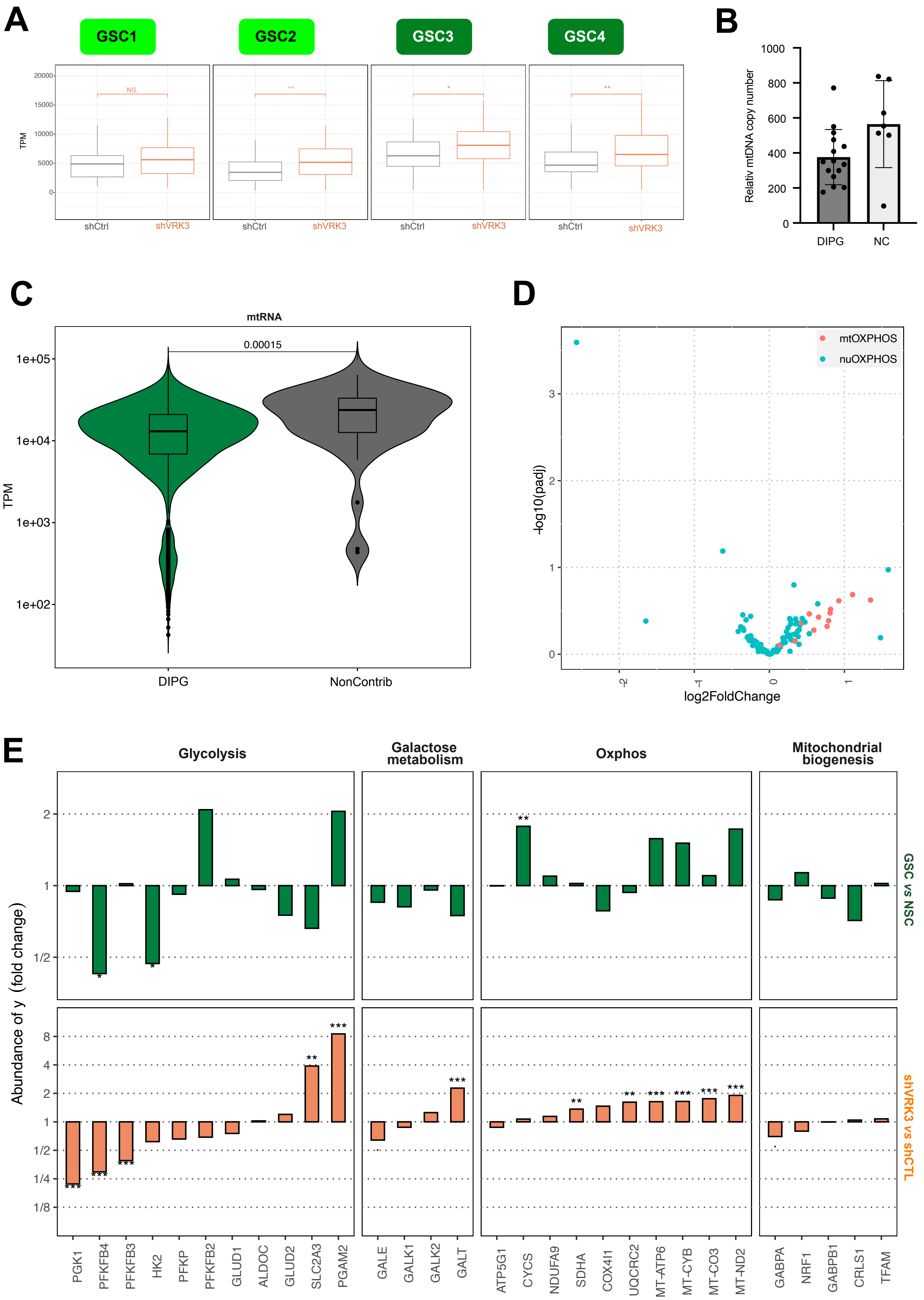
A
GSC1
GSC2
GSC3
GSC4
TPM
shCtrl
shVRK3
B
Relativ mtDNA copy number
DIPG
NC
C
mtRNA
0.00015
TPM
DIPG
NonContrib
D
mtOXPHOS
nuOXPHOS
-log10(padj)
log2FoldChange
E
Glycolysis
Galactose metabolism
Oxphos
Mitochondrial biogenesis
Abundance of y (fold change)
GSC vs NSC
shVRK3 vs shCTL
PGK1
PFKFB4
PFKFB3
HK2
PFKP
PFKFB2
GLUD1
ALDOC
GLUD2
SLC2A3
PGAM2
GALE
GALK1
GALK2
GALT
ATP5G1
CYCS
NDUFA9
SDHA
COX4I1
UQCRC2
MT-ATP6
MT-CYB
MT-CO3
MT-ND2
GABPA
NRF1
GABPB1
CRLS1
TFAM
